# Sulfur limitation increases duckweed starch accumulation without compromising growth

**DOI:** 10.1101/2021.02.22.432231

**Authors:** Zuoliang Sun, Wenjun Guo, Xuyao Zhao, Jingjing Yang, Pengfei Duan, Shuqing Xu, Hongwei Hou

**Author notes:** These authors contributed equally to this work (Zuoliang Sun, Wenjun Guo).

## Abstract

Duckweeds contain relatively high levels of starch and are a potential biomass feedstock for biofuel production. Here, the biomass and starch yield of duckweed under three different nutrient-limited conditions were analyzed to investigate possible ways of further increasing the efficiency of starch production. The results showed that sulfur limitation resulted in the highest starch yield, which was 42% and 73% higher than in nitrogen or phosphorus limitation, respectively. The high yield of sulfur-limited duckweed is largely due to the combinations of little effects on biomass and high accumulations of starch. Although nitrogen limitation led to higher starch content (67.4%), it severely reduced biomass production. The photosynthetic performance indicator *Fv*/*Fm* was a simple and sensitive indicator of starch content in nutrient-limited duckweed. Taken together, this study demonstrates that sulfur limitation is a simple and efficient way to increase starch yield, highlighting the great potential of duckweed for biofuel production. We report that sulfur limitation is a practical approach to increase starch yields in duckweed without affecting growth or biomass.

**Highlights:**

1. Sulfur limitation induces starch production in a duckweed specie.
2. Nitrogen limitation triggers the highest starch content, but limits growth.
3. Sulfur limitation results in the highest starch yield.
4. *Fv*/*Fm* is a rapid and robust proxy of starch content in nutrient-limited duckweed.

## 1. Introduction

The development of alternative fuels is becoming increasingly urgent as the world supply of fossil fuels decreases and atmospheric CO_2_ levels continue to rise. Microalgae and cyanobacteria have been identified as promising biological sources of various fuel-relevant molecules including lipids, ethanol, and hydrocarbons. As global CO_2_ levels rise and fossil fuel abundance decreases, the development of alternative energy becomes increasingly imperative (Kaur et al., 2017). Although bioethanol has emerged as a promising alternative energy source, it is constrained by economic fallout and effect on human food security (Papong and Malakul, 2010; Sanchez and Cardona, 2008), as the production of bioethanol is largely based on traditional land crops (such as maize [*Zea mays*], sweet potato [*Ipomoea batatas*], and cassava [*Manihot esculenta*]) on prized arable land at the detriment of foodstuffs. Therefore, it is an urgent need to identify novel and sustainable feedstock that is rich in starch for bioethanol production. Recently, duckweeds, a group of small floating aquatic plants (Appenroth et al., 2013), have gained much attention as a potential feedstock for bioenergy production due to their fast growth rate, low nutritional requirements, and high starch content (Cui and Cheng, 2015; Xu et al., 2011, 2012).

The biomass yield of duckweeds can be as high as 106 t dry weight (DW)/ha/year, which is much higher than that of the traditional crop wheat (*Triticum aestivum*), barley (*Hordeum vulgare*), or maize (Cui and Cheng, 2015). Depending on the species and growing conditions, duckweed starch content can range from 3 to 75% of its dry weight (Reid and Bieleski, 1970). Furthermore, duckweed starch can be easily converted into bioethanol via the existing fermentation method that is used for corn starch (Ge et al., 2012). Because duckweed biomass contains less lignin and cellulose than that of terrestrial plants, producing bioethanol from duckweed biomass does not require complex pre-treatments and is thus much more cost-effective (Cheng and Stomp, 2009; Ge et al., 2012). Notably, duckweed biomass is much easier to collect in large quantities compared to microalgae, which require energy-intensive and time-consuming centrifugation or membrane filtration steps for harvesting. Duckweeds also could be used to treat wastewater in wastewater ponds and then be collected for bioethanol production, which would alleviate land use pressures (Cheng and Stomp, 2009).

Biologically derived bioenergy from duckweed are promising sources, but improvements throughout the production process are required to reduce operational cost. Increasing starch yields in duckweed without compromising growth has great potential to improve economic feasibility. The high growth rates and starch yield are vital prerequisites for the large-scale application of duckweeds for bioenergy production. To this end, it is essential to optimize cultivation strategies to further enhance starch production while reducing the cost of duckweed bioprocessing. Starch is the main photosynthetic carbon sink in duckweed. Generally, duckweed contains starch at a low level when maintained under optimal growth conditions, but starch content drastically increases under stress conditions such as nutrient limitation, salinity, high light, phytohormone treatment, or shifting to a heterotrophic growth mode (Cheng and Stomp, 2009; Guo et al., 2020; Sun et al., 2020; Xu et al., 2012; Yin et al., 2015; Zhao et al., 2014; Zhou et al., 2018). The macronutrients nitrogen (N), phosphorus (P), and sulfur (S) are essential for plant growth and development, participating in the formation of cell structures and high energy molecules (nucleic acids, proteins, chlorophyll, ATP, and phospholipids) (Cai et al., 2013; Sun et al., 2013). Duckweed can survive and even proliferate in the absence of these macronutrients for a long period of time (Cheng and Stomp, 2009). Currently, the most intensively studied starch induction methods in duckweed are N limitation (NL) and P limitation (PL) (Guo et al., 2020; Reid and Bieleski, 1970; Tao et al., 2013; Zhao et al. 2015). However, these two strategies trigger starch accumulation in duckweed at the expense of biomass production. Hence, despite a higher starch content, starch yield may be significantly compromised due to the tradeoff between starch induction and plant growth. In contrast, much less attention has been paid to the effects of sulfur limitation (SL) in the context of growth and starch accumulation in duckweed. As with microalgae, mounting evidence suggests that SL could be a more efficient strategy than NL or PL to stimulate strong starch or lipid production (Mao et al., 2018; Vitova et al., 2015; Yao et al., 2012). As such, a systematic study of the effects of SL on growth and starch accumulation in duckweed is needed.

Here, a strain of the giant duckweed, *Spirodela polyrhiza*, which is a suitable candidate in the Hg bioremediation system (Yang et al., 2018), was focused on. Its growth characteristics and starch accumulation in response to various nutrient limitations (SL, PL, and NL) were investigated. In addition, the comparative analysis on chlorophyll and protein contents, photosynthetic efficiency, as determined by *Fv*/*Fm* measurements, and Rubisco activity among different growth conditions also were performed. This study provides a biochemical and physiological framework for duckweed-based starch biosynthesis and its application to bioenergy production.

## 2. Materials and Methods

### 2.1 Duckweed strain and culture conditions

The axenic strain *Spirodela polyrhiza* (L.) Schleid (5543) was used, isolated from East Lake (N 30°32′, E 114°21′), Wuhan, China, in September 2014. A culture was maintained in 600 mL half-strength Murashige and Skoog (MS) medium, pH 5.8, and supplemented with 10 g/L glucose as a propagation medium. Cultures were kept at 25°C for 7 days and used for subsequent experiments, under a light intensity of 5,000 lux and a 16 h light:8 h dark photoperiod.

After 7 days of cultivation, plants were harvested and washed in a growth medium lacking either sulfur (S), nitrogen (N), or phosphorus (P). The remaining traces of medium were removed by patting the plants dry on paper towels before transferring plants to flasks containing S-, N-, or P-free growth medium. For P limitation (PL) or N limitation (NL), P (in the form of KH_2_PO_4_) or N (in the form of KNO_3_ and NH_4_NO_3_) was omitted from the medium; KH_2_PO_4_ and KNO_3_ were replaced with KCl; for S limitation (SL), MnSO_4_, MgSO_4_, CuSO_4_, and ZnSO_4_ in the growth medium were replaced with MnCl_2_, MgCl_2_, CuCl_2_, and ZnCl_2_, respectively, in matching concentrations. In all cases, the desired nutrient limitation in the modified growth medium was confirmed by chemical analysis. All cultures were grown in 1 L glass beakers with a working volume of 600 mL (radius: 5.5 cm) of half-strength MS medium supplemented with 10 g/L glucose, which was inoculated with 2 g of actively growing plants to cover the entire surface of the liquid medium with a single layer of fronds. Half-strength MS medium with 10 g/L glucose as organic carbon source was used as control. Beakers were maintained at 25°C under a 16 h light: 8 h dark photoperiod with a light intensity of 5,000 lux for 14 days. All treatments were performed in triplicates. Duckweed plant tissue was collected at 0.5, 1, 4, 7, 10, and 14 days after the beginning of treatment, using the entire beaker as one sample for physiological and biochemical analysis.

### 2.2 Determination of pigment contents

The chlorophyll content was determined as previously published described (Liu et al., 2019). Briefly, 0.05 g of fresh plant tissue was homogenized in 1 mL of 95% (v/v) ethanol and kept at 4°C for 48 h in the dark. Extracts were then centrifuged at 10,000 × *g* for 5 min at 4°C. The supernatant was used for spectrophotometric determination of total chlorophyll content.

### 2.3 Growth measurements

Fresh duckweed was collected and patted dry on paper towels for 5 min, after which fresh weight was determined using a precision balance (Sartorius, Germany) (Bergmann et al., 2000). Then plant tissues were dried at 60°C until the weight stayed constant, at which time dry weight was measured. The growth rate was calculated according to the formula Growth rate = increased dry weight/area/time (g/m^2^/day).

### 2.4 Determination of starch content

Starch content was determined using a Starch Assay Kit (Suzhou Grace Bio-technology Co., Ltd., Suzhou, China). Briefly, dried plant tissues were ground into powder using a mortar and pestle. Then 10 mg powder was homogenized with 0.5 mL Dimethyl sulfoxide (DMSO). The homogenates were heated in a boiling water bath until completely dissolved. Samples were subsequently allowed to cool down to room temperature, and 1.5 mL ethanol was added, followed by mixing by vortexing and centrifugation at 10,000 × *g* for 5 min at 25°C. The supernatant was discarded and the remaining pellet was air-dried before being resuspended with 0.5 mL DMSO and boiled for 15 min. The suspension was centrifuged again at 10,000 × *g* for 5 min at 25°C and the supernatant was used as samples for starch assays according to the manufacturer’s instructions. The starch yield was calculated with the formula: starch yield (g/m^2^) = biomass yield × starch content.

### 2.5 Determination of chlorophyll fluorescence

To determine chlorophyll fluorescence, duckweed fronds were collected from each growth medium and dark-adapted for 10 min. Then chlorophyll fluorescence was measured with a plant efficiency analyzer (PEA, Hansatech Ltd, UK). During measurements, plants were first exposed to low irradiation to assess basal chlorophyll fluorescence (*F_0_*). A saturating light flash was then applied to determine maximum fluorescence (*F_m_*). The maximum efficiency of photosystem II (PSII) *Fv*/*Fm* was calculated following the equation: *Fv*/*Fm* = (*Fm-F_0_*)/*Fm* and indicates the maximal quantum yield of PS II photochemistry, reflecting the maximal light energy conversion efficiency of PS II (Baker, 2008).

### 2.6 Analysis of protein content and photosynthetic enzyme activity

Freshly harvested plant tissue (0.1 g) was homogenized in 0.9 mL of 1x phosphate-buffered saline (PBS, pH 7.4) in an ice water bath. The extracts were then centrifuged at 10,000 × *g* for 10 min at 4°C. Total protein content was determined in the supernatants using the Protein Assay Kit (Nanjing Jiancheng Bioengineering Co., Ltd., Nanjing, China) following the manufacturer’s instructions.

To assay the activity of enzymes, 0.1 g fresh duckweed was homogenized in an ice-cold mortar with 1 mL of 8 mM MgCl_2_, 2 mM EDTA, 50 mM HEPES-NaOH pH=7.6, 5 mM DL-Dithiothreitol, 2% (w/v) polyvinylpyrrolidone-40, and 12.5% (w/v) glycerol. The homogenate was then centrifuged at 10,000 × *g* for 5 min at 4°C. The supernatant was used to measure enzyme activity. Rubisco activity was determined using spectrophotometric diagnostic kits (Suzhou Grace Bio-technology Co., Ltd., Suzhou, China).

### 2.7 Statistical analysis

All results are shown as mean values ± standard deviation of three replicates for each treatment. One-way ANOVA was performed to compare all data. In both cases, differences between the analyzed variables were considered significant according to the LSD test (*P* < 0.05). All analyses were performed using SPSS 19.0 (SPSS, Chicago, IL, USA).

## 3. Results and Discussion

### 3.1 Growth characteristics and chlorophyll content

Nutrients are essential elements for plant growth and development. Nutrient limitations can trigger morphological and physiological responses in plants. When the giant duckweed grew in NL, PL, and SL conditions, clear morphological changes were found (Supplemental Fig. S1 and S2). The plants showed a morphological response to NL as early as 4 days after the beginning of the treatment, with SL fronds only exhibiting gross morphology changes after 7-10 days and P-limited plants taking a full 14 days to develop a visual phenotypic response (Supplemental Fig. S2). Specifically, NL and SL both resulted in yellow fronds due to a rapid reduction in total chlorophyll content (Fig. 1), in addition to a higher root-to-shoot ratio, bending of the fronds, and anthocyanins accumulation. P-limited plants had dark green leaves, which may be attributed to their higher chlorophyll levels (Fig. 1). The present results are in agreement with Reid and Bieleski (1970), who demonstrated that newly emerged duckweed leaves were dark green and that the plants had elongated roots in P-limited conditions. Similar phenotypes have also been observed in the context of other nutritional stresses, such as N, P, or magnesium limitation in other species (Hermans et al., 2006; Rao and Terry, 1989; Zhao et al., 2015). Nutrient limitations generally result in more carbon being allocated to the root to support more root growth and increase the uptake of these nutrients. They therefore affect sugar metabolism and/or carbohydrate partitioning between source and sink tissues, and thus photosynthesis performance (Hermans et al., 2006).

**Fig. 1.**
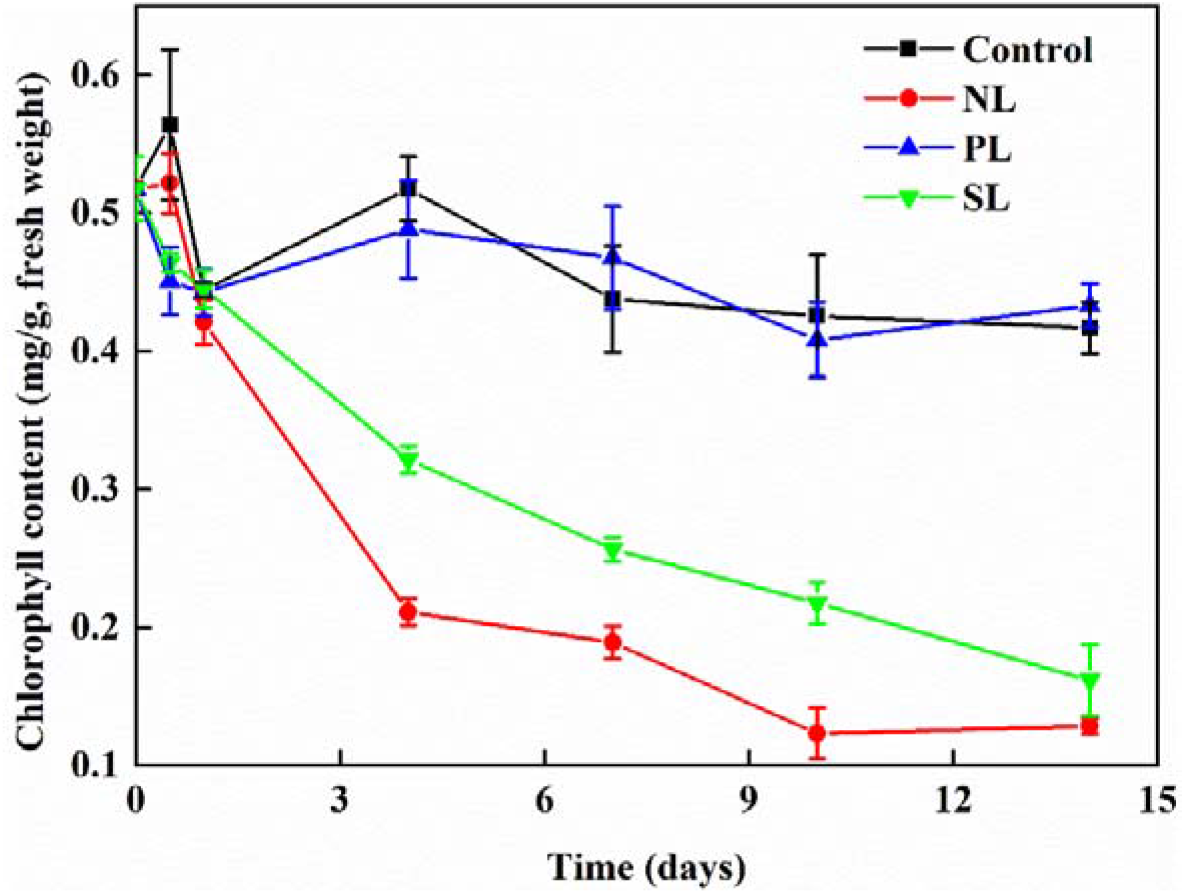
Chlorophyll content of duckweed grown in various nutrient limitation conditions for 14 days. NL: nitrogen limitation; PL: phosphorus limitation; SL: sulfur limitation.

Nutrient limitation also can adversely affect chlorophyll levels (Daughtry et al., 2000; Zhao et al., 2003). Chlorophyll content diminished when duckweed was grown in SL or NL conditions, but not under PL (Fig. 1). NL initiated a rapid drop in chlorophyll content, reaching only 24.9% of their initial values after 14 days of cultivation. Similarly, chlorophyll content at the end of cultivation in SL conditions was 31.2% of initial values. Finally, plants growing in PL conditions did not show statistical difference in chlorophyll content in comparison to the one growing in control conditions. These results broadly agreed with morphological observations (Supplemental Fig. S2), and also suggested that nutrient limitations might attenuate the photosynthetic performance of duckweed. Similarly, Liu et al. (2018) showed that nutrient limitation (imposed as distilled water) negatively regulated total chlorophyll biosynthesis and net photosynthetic rates. Tao et al. (2013) also suggested that nutrient limitation significantly inhibited photosynthesis.

Limitations of N, P or S in medium also altered growth rates in the giant duckweed (Fig. 2). Plants grown under NL or PL had lower dry weight in comparison to the control group, with a decrease of 18% (PL) and 42% (NL). Surprisingly, the omission of sulfur from the growth medium had no detrimental consequences on dry weight (Fig. 2). After growth for 14 days, the dry weight of plants growing under SL was thus 81% higher than in NL conditions and 29% higher than in PL conditions. The relatively little effects of SL on dry weight growth rates of the duckweeds are likely due to the fact that the duckweed plants have abundant endogenous sulfur stores that may fulfill the S quota necessary to support normal growth for the 14 days covered by this study. Ferreira and Teixeira (1992) reported that the duckweed *Lemna minor* continued to divide and remain viable for at least 26 days when cultured in growth medium lacking sulfur. In addition, as is the case for other plants, the amount of sulfur duckweed needs to sustain normal plant growth (the S quota) is much lower than that for P and N (Epstein, 1972). So that SL is not associated with phenotypes as severe as those seen during PL or NL. Previous reports have demonstrated that nutrient deficiency (N, P) can severely and negatively influence duckweed biomass yield (Reid and Bieleski, 1970; Zhao et al., 2015). Guo et al. (2020) reported a dry weight of 5.6 g/m^2^/day under NL for the duckweed *Landoltia punctata* 0202, which is even lower than the dry weight measured under similar conditions here. Similarly, Liu et al. (2018) measured a dry weight of 5.6 g/m^2^/day after 14 days of cultivation for *L. punctata* 0202 in distilled water. The various biomass yields obtained under nutrient limitation conditions are summarized in Table 1. After 14 days of cultivation, duckweed grown in SL conditions therefore accumulated 26% and 71% more dry biomass than those grown in PL or NL conditions, respectively.

**Fig. 2.**
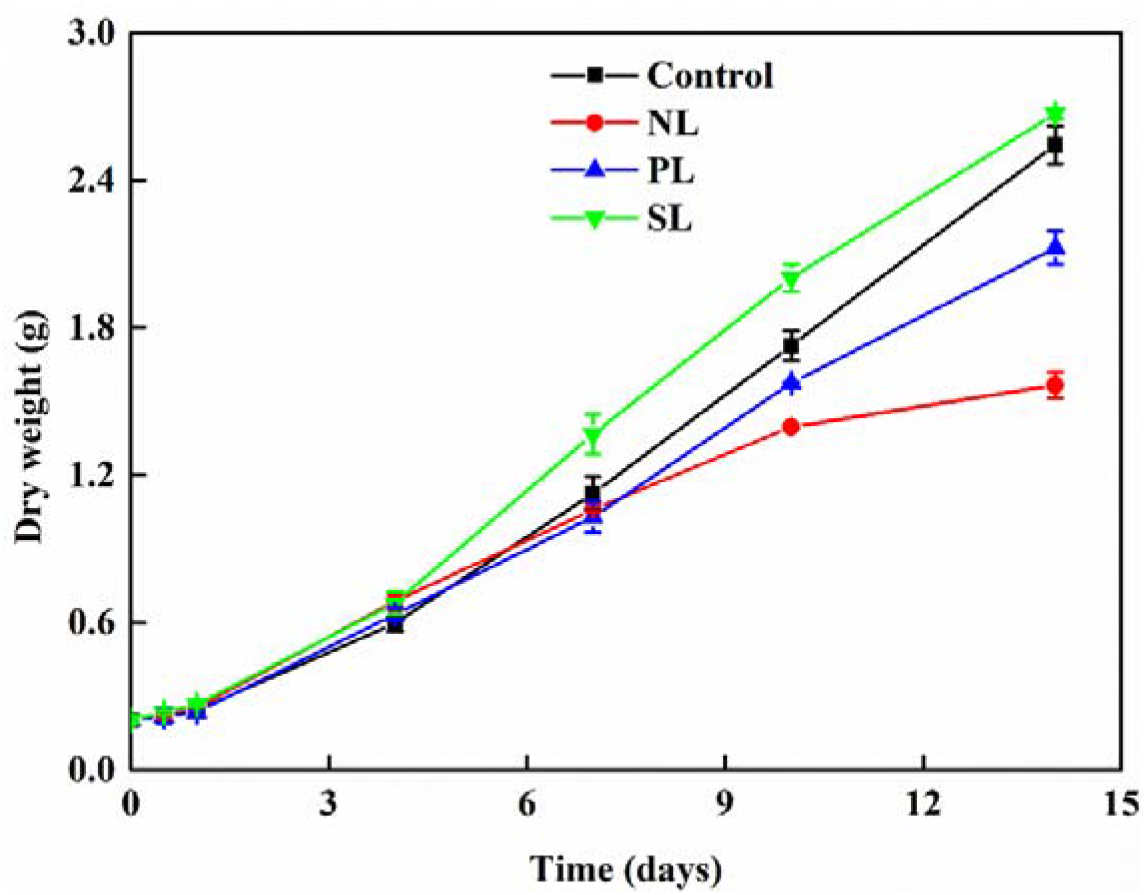
Growth parameters of duckweed under different nutrient limitation conditions. NL: nitrogen limitation; PL: phosphorus limitation; SL: sulfur limitation.

**Table 1.** Duckweed growth parameters under different nutrient limitation conditions after 14 days of cultivation. C: control; SL: sulfur limitation; NL: nitrogen limitation; PL: phosphorus limitation.

| Cultivation conditions | Biomass yield<br>(g/m<sup>2</sup>) | Starch content<br>(%, dry weight) | Starch yield<br>(g/m<sup>2</sup>) | Starch productivity<br>(g/m<sup>2</sup>/day) |
| --- | --- | --- | --- | --- |
| C | 258.2 ± 7.7 a | 24.5 ± 0.6 d | 63.3 ± 7.7 d | 4.5 ± 0.5 d |
| SL | 271.1 ± 2.0 a | 56.2 ± 1.2 b | 152.3 ± 2.1 a | 10.9 ± 0.2 a |
| NL | 159.0 ± 5.4 c | 67.4 ± 3.4 a | 107.2 ± 5.1 b | 7.7 ± 0.4 b |
| PL | 215.6 ± 6.8 b | 40.6 ± 0.9 c | 87.6 ± 2.2 c | 6.3 ± 0.1 c |

Different lowercase letters in the same column indicate significant differences according to LSD test ( $P < 0.05$ ). Values represent mean ± SD ( $n=3$ ).

### 3.2 Starch content and yield under various stress conditions

To serve as an alternative feedstock for biofuel production, the starch yield, which is determined by biomass and starch content, is critical. Then the starch content in the duckweed plants cultivated in SL, PL, and N conditions was quantified. Starch content gradually increased over the 14 days of growth under all three nutrient limitation conditions (Fig. 3a). When measured by dry weight, after 14 days, the starch content of the plants grew under control conditions was 24.5%, whereas the starch content reached 40.6% for plants grew under PL conditions, 56.2% under SL conditions, and 67.4% under NL conditions (Fig. 3a). Zhao et al. (2015) reported that duckweed starch content increased from 8.9% in replete medium to 32.5% and 23.0% after 15 days of cultivation in NL or PL conditions, respectively, which is consistent with the present results that NL is more efficient than PL in inducing starch accumulation. The currently highest reported starch content for duckweed was achieved after 9 days of cultivation in NL conditions in the presence of sucrose, reaching 60.0% of the total dry weight (Yu et al., 2017), a value that is comparable to the results with NL conditions here. Guo et al. (2020) also reported that duckweed can quickly accumulate starch up to 52.4% under NL. Similarly, by transferring *S. polyrhiza* from nutrient-replete conditions to tap water for 5 days, the starch content increased from about 20.0% to 45.8% of total dry weight (Cheng and Stomp, 2009). The maximum starch content for duckweed measured here (NL: 67.4%; SL: 56.2%) reached roughly the same levels reported in barley (51.3 ~ 64.2%) (Guo et al., 2020), indicating that starch-enriched duckweed could be regarded as a potential feedstock for starch production.

**Fig. 3.**
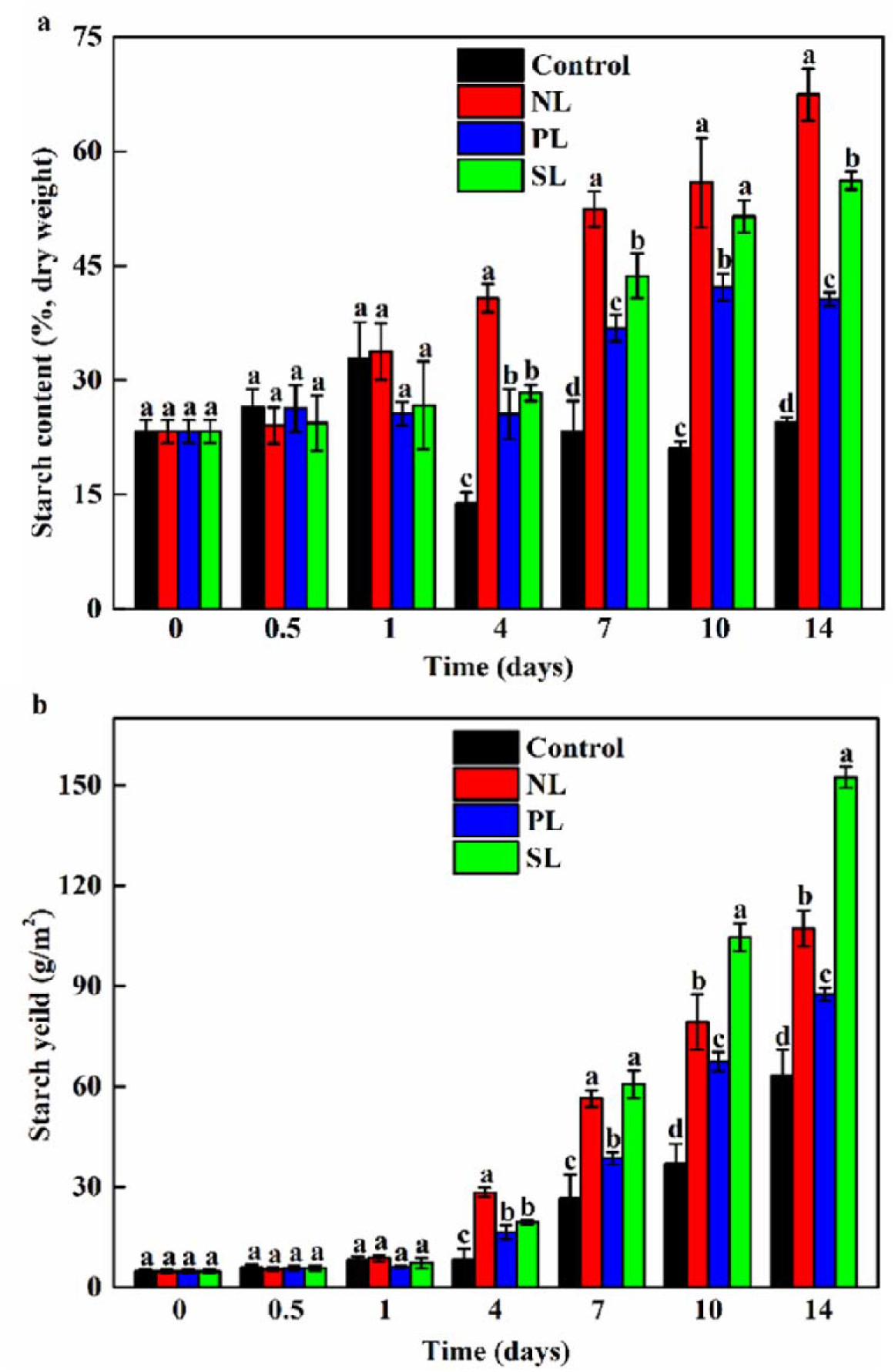
Starch content and yield of duckweed grown in various nutrient limitation conditions for 14 days. (a) starch content; (b) starch yield. NL: nitrogen limitation; PL: phosphorus limitation; SL: sulfur limitation. Different lowercase letters indicate significant differences according to the LSD test (*P* < 0.05).

Based on the starch content and biomass information, the starch yield was estimated under different growth conditions (Table 1 and Fig. 3b). Under all three nutrient limitation conditions, the giant duckweed increased starch yield in comparison to the control group. The highest yield was found in plants grown under SL conditions (Fig. 3b). Starch yield of S-limited duckweed increased sharply over time. Although the yield was initially higher than in P-limited plants but lower than N-limited plants, starch yield in SL growth conditions surpassed that obtained with NL after 10 days of cultivation. Extending the cultivation time saw a further rise in starch yield (Fig. 3b). Duckweed grown in SL conditions accumulated 42% and 73% more starch than when grown in NL or PL conditions, respectively (Table 1). The mean starch productivity achieved 10.9 g/m^2^/day under SL conditions, which is 1.4 times higher than the control group (Table 1). The observed high starch yield is also higher than previous studies in other duckweed species. For example, Guo et al. (2020) reported a starch productivity of 3.9 g/m^2^/day for *Landoltia punctata* grown under NL. Zhao et al. (2015) obtained a starch productivity value of 1.8 g/m^2^/day and 1.0 g/m^2^/day in NL and PL conditions, respectively. The nutrient limitation trigged starch yield increase was also found in an industrial-scaled *S. polyrhiza* growing system. By pre-cultivating duckweed in diluted pig effluent before transfer into clean water for 10 days of cultivation, the starch content in *S. polyrhiza* increased by 64.9% compared to the control group, and the productivity reached 2.6 g/m^2^/day (Liu et al., 2020), likely due to the limitation of several nutrients (e.g., N, P and S). As this study showed that NL and PL reduced biomass production significantly, further optimizing the industrial *S. polyrhiza* growing system by specifically subjecting plants to SL could further increase the starch productivity to the level equal or higher than in maize Guo et al. (2020).

### 3.3 Photosynthetic activity

*Fv*/*Fm* measures the maximum quantum efficiency of photosynthetic tissues and is thus an important indicator of plant photosynthetic activity. *Fv*/*Fm* also provides a sensitive and intrinsic indicator of stress conditions: healthy plants have values > 0.8 (Lan et al., 2010); however, *Fv*/*Fm* will decrease in photosynthetic organisms suffering from some environmental stress (Baker, 2008; Lan et al., 2010).

To explore how nutrient limitation affects photosynthesis performance in duckweed, *Fv*/*Fm* therefore was measured under NL, PL, and SL conditions. The initial value of *Fv*/*Fm* was 0.84 in duckweed from the control group grown in a replete growing medium. *Fv*/*Fm* declined in the three nutrient limitation conditions compared to the control group (Fig. 4), indicative of lower photosynthetic activity. PL had a much milder effect on *Fv*/*Fm* values relative to the other two nutrient limitations, as *Fv*/*Fm* values after 14 days of cultivation in PL conditions remained fairly high, having dropped from 0.84 to 0.69. In contrast, *Fv*/*Fm* exhibited a rapid response to NL conditions, with a reduction in values from 0.84 to 0.30 by day 14. SL caused an intermediate effect on *Fv*/*Fm*, with final values after 14 days of SL around 0.52. These results demonstrated that NL resulted in a more severe loss of photosystem II (PSII) activity than either SL or PL conditions. When comparing the results presented in Fig. 3a with Fig. 4, the increase in starch content paralleled the decrease in *Fv*/*Fm* was noticed, with lower *Fv*/*Fm* values obtained from plants with higher starch content. Therefore, *Fv*/*Fm* may be regarded as a convenient, although indirect, indicator of starch content.

**Fig. 4.**
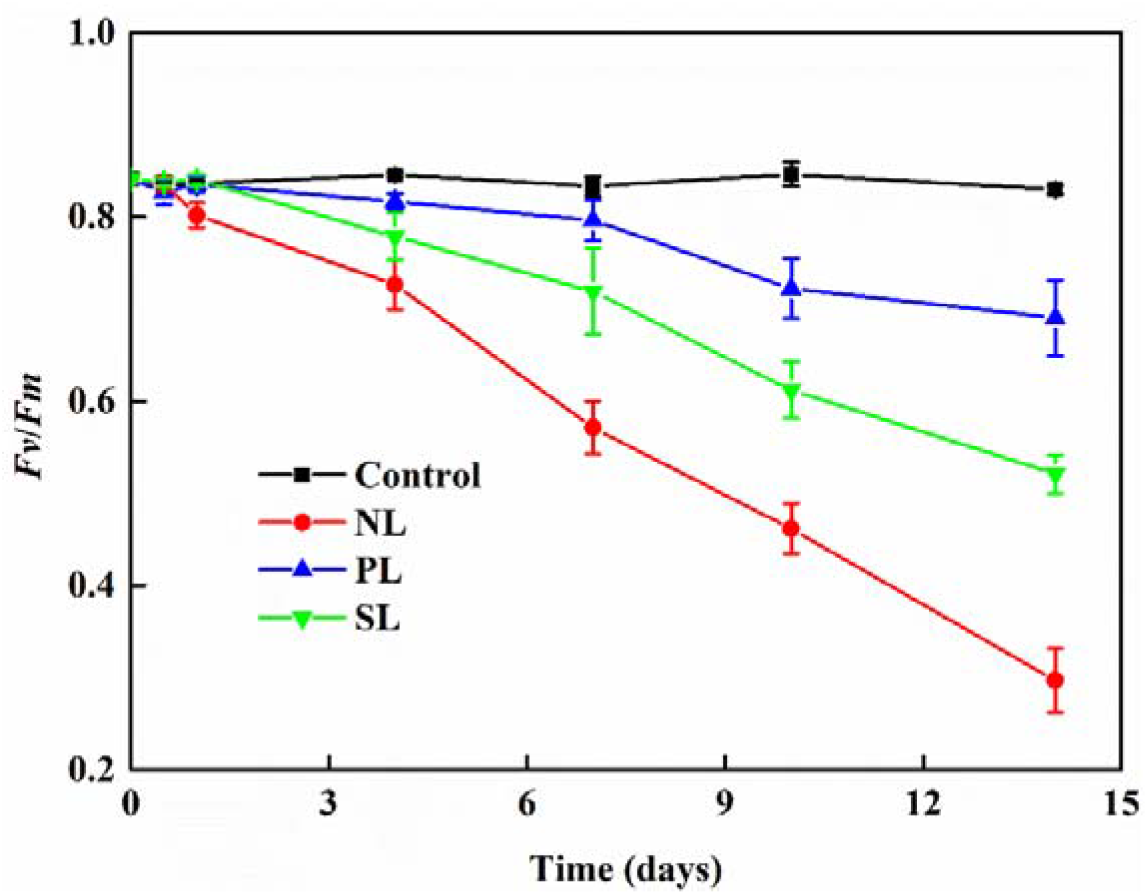
*Fv*/*Fm* of duckweed grown in various nutrient limitation conditions for 14 days. NL: nitrogen limitation; PL: phosphorus limitation; SL: sulfur limitation.

### 3.4 Protein content and Rubisco activity

Protein content initially stayed constant in N-limited plants during the first 4 days of treatment but then progressively declined over the course of the following 10 days (Fig. 5a). Plants exposed to PL and SL already displayed lower protein content by day 4, and protein content only decreased more as time progressed. By the end of the 14 days of cultivation, total protein content was 64.1%, 65.6%, and 50.4% of initial values in NL, PL, and SL conditions, respectively. Previous work reported that nutrient limitation (for N or P) leads to protein degradation in duckweed, as a means of recycling amino acids (Zhao et al., 2015), which is in line with the findings here. Based on these results, protein and starch contents were proposed to be inversely correlated under nutrient limitation conditions (Zhao et al., 2015). Carbohydrates, lipids, and protein are vital building blocks for maintaining normal plant function under adverse conditions like nutrient limitation. Duckweed appears to allocate its energy stores to starch production while degrading proteins under nutrient limitation. Chlorophyll and protein biosynthesis are both blocked under nutrient scarcity; in addition, the flow of carbon is rerouted to starch in duckweed. These findings in this work revealed a reassignment of metabolic carbon flow from proteins toward starch biosynthesis in nutrient-limited duckweed.

**Fig. 5.**
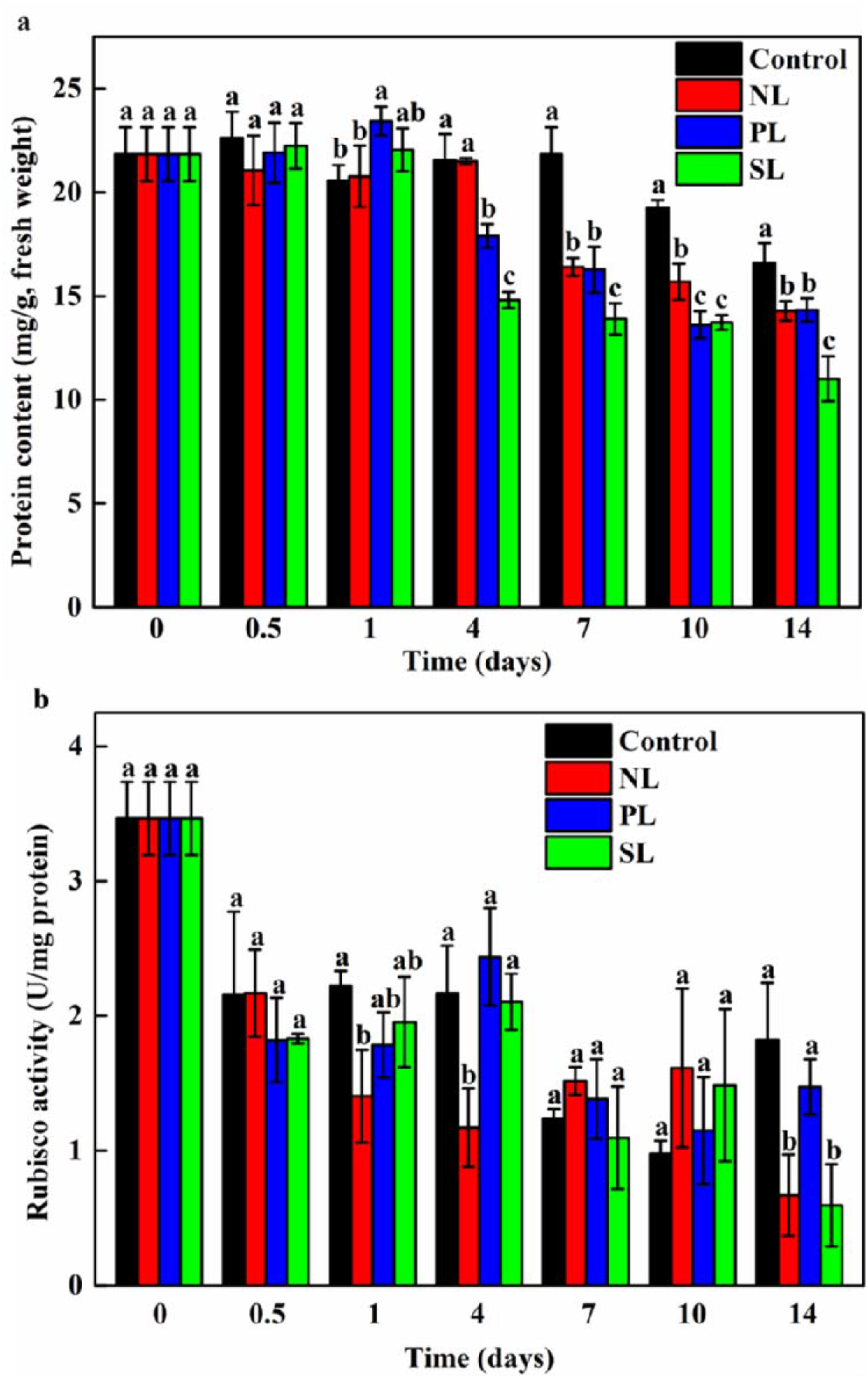
Protein content and Rubisco activity in response to different stress conditions during duckweed cultivation. (a) protein content; (b) Rubisco activity. NL: nitrogen limitation; PL: phosphorus limitation; SL: sulfur limitation. Different lowercase letters indicate significant differences according to the LSD test (*P* < 0.05).

To elucidate the effects of nutrient limitation on photosynthetic performance, Rubisco activity was measured. After 14 days of cultivation, Rubisco activity globally decreased over time in both the control and test groups (Fig. 5b). Closer inspection revealed that Rubisco activity in P-limited plants was largely comparable to that in the untreated control group. In contrast, both N- and S-limited plants had lower Rubisco activity relative to the control group at the end of the cultivation period. These results validated the measurements of *Fv*/*Fm* and chlorophyll that suggested that Rubisco was not sensitive to PL.

## 4. Conclusions

Duckweed induced starch accumulation under different nutrient limitation conditions, accompanied by a significant downregulation of Rubisco activity, photosynthetic efficiency (*Fv*/*Fm*), and chlorophyll and protein content. The highest starch content and highest starch productivity were obtained under NL (67.4%, dry weight) and SL (10.9 g/m^2^/day) conditions, respectively. Consequently, SL may provide a promising new avenue for boosting starch accumulation in duckweed for various biotechnological applications.

## Acknowledgements

This work was supported by the Major Project of Technology Innovation Program of Hubei, China (2017ABA135); the National Key R & D Program of China (2018YFD0900801); Open Project of Guangdong Provincial Key Laboratory of Applied Botany, South China Botanical Garden, Chinese Academy of Sciences (AB2018020); and Project of State Key Laboratory of Freshwater Ecology and Biotechnology (2019FB11). The authors would like to thank Xin Wang at the Analysis and Testing Center of Institute of Hydrobiology, Chinese Academy of Sciences for experimental support.

